# Digenic variant interpretation with hypothesis-driven explainable AI

**DOI:** 10.1101/2023.10.02.560464

**Authors:** Federica De Paoli, Giovanna Nicora, Silvia Berardelli, Andrea Gazzo, Riccardo Bellazzi, Paolo Magni, Ettore Rizzo, Ivan Limongelli, Susanna Zucca

## Abstract

**Motivation:** The digenic inheritance hypothesis holds the potential to enhance diagnostic yield in rare diseases. Computational approaches capable of accurately interpreting and prioritizing digenic combinations based on the proband’s phenotypic profiles and familial information can provide valuable assistance to clinicians during the diagnostic process.

**Results:** We have developed diVas, a hypothesis-driven machine learning approach that can effectively interpret genomic variants across different gene pairs. DiVas demonstrates strong performance both in classifying and prioritizing causative pairs, consistently placing them within the top positions across 11 real cases (achieving 73% sensitivity and a median ranking of 3). Additionally, diVas exploits Explainable Artificial Intelligence (XAI) to dissect the digenic disease mechanism for predicted positive pairs.

**Availability and Implementation:** Prediction results of the diVas method on a high-confidence, comprehensive, manually curated dataset of known digenic combinations are available at oliver.engenome.com.

## 1. Introduction

The identification of disease-causing mutations represents a paramount challenge in the realm of human genetics. Rare Diseases (RDs), which affect around 6% of the population in Western societies, display a genetic component in roughly 80% of instances^1^. Precise genetic diagnosis not only leads to better disease management but also aids in tailoring the most appropriate treatment strategy for each patient^2^. Recent years have witnessed remarkable advancements in Whole Genome Sequencing (WGS) and Whole Exome Sequencing (WES), enhancing our capacity to uncover the genetic underpinnings of RDs^3–5^ and employing computational tools that facilitate variant interpretation and prioritization has proven to be of paramount importance^6^. In the case of individuals suspected of having a RD, the diagnostic process aims to identify the potential genetic cause by considering the patient’s phenotypic characteristics and familial information. Predominantly, most RDs exhibit monogenic (Mendelian) inheritance^7^, prompting clinicians to focus on identifying the single mutated gene responsible for the observed disease phenotype(s).

To achieve this, the American College of Medical Genetics (ACMG), in collaboration with the Association for Molecular Pathology (AMP), has formulated guidelines that aid in the interpretation of genomic variants as pathogenic, likely pathogenic, benign, likely benign, or of uncertain significance (VUS). These determinations are based on diverse factors, including familial segregation, *in silico* predictions of the variant’s damaging impact, and population allele frequency^8^. Medical professionals employ these guidelines to identify the potentially small number of pathogenic variants in a patient, ultimately aiming to pinpoint the mutation causing the disease.

Despite the advancement in technical capabilities and knowledge, the diagnostic yield, defined as the percentage of patients for whom the genetic cause of the disease is identified after sequencing, varies between 35% to 55% based on the disorder^1^, leaving approximately 200 million patients without a definitive diagnosis^5^.

In contrast to the monogenic inheritance hypothesis, oligogenic models propose that the manifestation of a genetic disorder arises from the combined occurrence of mutations in multiple genes. The simplest form of this paradigm is digenic inheritance (DI), wherein the disease’s origin is attributed to two mutated genes. In the DI field, distinct mechanisms have been recognized: True Digenic (TD, also known as Pure Digenic), Composite (CO, also known as Modifier), or Dual Molecular Diagnosis (DM). In the case of TD, the disease’s phenotype(s) emerges when both causative genes carry mutations in the patient’s genome. Modifiers denote variants that, when present alongside causative variants, can influence disease outcomes, such as severity or onset age. Lastly, the DM mechanism entails the co-occurrence of two distinct disorders, each caused by pathogenic mutations in different genes^9^. The DI paradigm holds significant potential for supporting the diagnosis of RDs^7^. For instance, research has revealed various genes acting as digenic contributors in arrhythmogenic cardiomyopathy, a disorder typically regarded as monogenic^10^. Several other conditions have also been investigated in the context of DI patterns, including hypogonadotropic hypogonadism, deafness, ciliopathies, and Long QT syndrome^11^.

To pinpoint causative gene pairs, potentially thousands of candidates from WGS/WES must be assessed. Computational tools, exploiting statistical and Machine Learning (ML) techniques, greatly support the diagnostic procedure. Recent years have seen the emergence of numerous tools that assess the impact of individual mutations^12–14^, or predict variants’ pathogenicity^15–17^. Some methods further incorporate the patient’s phenotypic data to prioritize variants, aiming to discern the causative one among the top contenders^18^. A limited number of computational tools have been specifically designed to classify and prioritize digenic pairs^19^. VarCopp was the first published ML tool to categorize variant pairs as either pathogenic or benign, relying on 11 features grouped into two subsets: variant-level features (*in silico* predictions of the damaging effects of the variants in a pair) and gene-level features that capture both gene-gene interactions and *a priori* gene properties^20^. Subsequently, an additional ML model named Digenic Effect (DE) predictor was developed to further elucidate digenic mechanisms (TD, CO, or DM)^21^. ORVAL is an online platform that processes standard VCF files containing the proband’s variants, utilizing both VarCopp and the DE predictor for digenic classification and oligogenic network analysis^22^. DiGePred is a ML approach, available as a stand-alone tool. Like ORVAL, DiGePred uses different gene-gene interaction features, such as co-expression, protein-protein interaction and pathway similarity, and *a priori* gene-level features such as haploinsufficiency and gene essentiality. While DiGePred autonomously classifies gene pairs^23^, it doesn’t provide the variant-specific resolution that ORVAL offers. A more recent ML methodology for predicting digenic interactions at the gene level is DIEP (Digenic Interaction Effect Predictor)^24^. OligoPVP is an automatic variant interpretation tool that can be used to prioritize digenic and oligogenic instances^25^. To our understanding, OligoPVP is the sole tool that integrates proband’s phenotypes for interpretation. However, OligoPVP initially prioritizes each variant based on a pathogenicity score, subsequently prioritizing sets of two or more interacting genes as determined by a high-confidence gene-gene interaction network. The final score is the sum of the pathogenicity scores of all the variants involved in the combination. ORVAL, DiGePred, and DIEP were trained using known positive digenic pairs from DIDA^26^, a curated dataset listing 258 pathogenic variant pairs sourced from literature. Presently, DIDA is not available anymore, and the authors have recently published a new database, OLIDA, encompassing more than 1000 oligogenic variant combinations^27^. The need for expanded open-access resources concerning digenic and oligogenic variants, similar to ClinVar^28^, is crucial for understanding these complex genetic interactions and for providing a valuable reference for clinicians and researchers, aiding in accurate diagnoses and treatment decisions. Additionally, it’s essential to raise awareness and educate the community about digenic and oligogenic inheritance to promote collaboration and research progress. For this reason, there’s a pressing need to establish a community platform where researchers can share candidate causative combinations.

We introduce diVas, an ML tool designed for digenic variant interpretation, grounded in monogenic ACMG/AMP standard guidelines and gene-gene interaction data. DiVas can also harness the proband’s phenotypic data to enhance the classification and prioritization of digenic pairs. For positively predicted pairs, diVas employs cutting-edge Explainable Artificial Intelligence (XAI) techniques for further subclassification into distinct digenic mechanisms (TD/CO and DM). We have validated diVAs’ capability to identify the true causative pair from thousands of candidates in various real-world scenarios and have compared our tool with existing methods for digenic variant interpretation.

We applied this method to analyze our comprehensive high-confidence, manually curated dataset of digenic combinations and associated phenotypes. Predictions are available at https://oliver.engenome.com.

## 2. Materials and methods

### 2.1 Dataset description

#### 2.1.1 Training data collection and preprocessing

To collect data for training purposes, we curated a dataset of pathogenic digenic combinations extracted both from DIDA^26^, from literature review^29–32^ and from an internal unpublished database. Variants have been reported with their genomic coordinates and associated with the caused phenotypes expressed as Human Phenotype Ontology (HPO) terms^33^, if explicitly available. The positive training set contains 238 unique gene pairs and 316 unique genes. At the variant level, the distribution of variant effects in the training set is shown in Figure 1, with more than 90% having a coding effect. Taking into account both variants and HPO terms, each digenic combination in the training set is unique. For digenic pairs in the DIDA database that lacked HPO terms, we utilized phenoBERT^34^ to extract phenotypes from associated publications. In cases where scientific publications were unavailable, we employed a prevalence-based phenotype sampling method, drawing from the pool of phenotypes linked to the respective disease(s) in the HPO resource: the more frequently a phenotype appears in the manifestation of a disease, the more likely it is to be extracted when describing the digenic combination. Each causative digenic combination is associated with a median number of 5 HPO terms (range [1, 14]).

**Figure 1:**
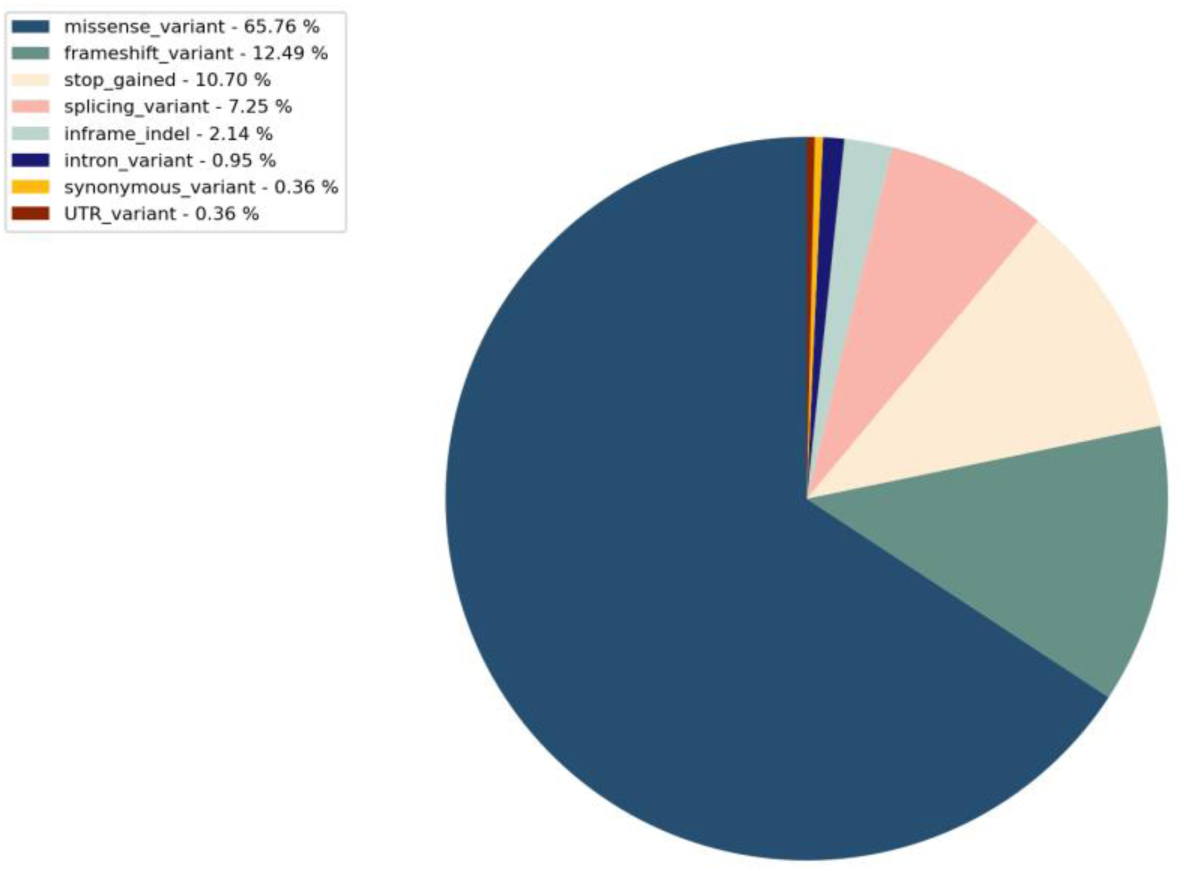
Distribution of Variant Effects in Pathogenic Digenic Combinations.

The negative dataset contains about 15,000 neutral digenic combinations extracted from 1000 Genomes Project (1KGP) data, and from solved monogenic or digenic real cases (internal data). Digenic combinations derived from the 1KGP were paired with either a randomly sampled set of phenotypes from a DIDA-positive digenic combination or a pool of phenotypes chosen through a prevalence-based approach from randomly selected disorders. For each real case, we first processed the VCF files to produce all potential digenic combinations. Subsequently, we selected a subset of these combinations that included:

i. at least one variant with a significant pathogenic impact on the gene (7% of the subset);
ii. one of the variants from the causative pair (3% of the subset); (iii) combinations chosen at random (90% of the subset). In summary, pathogenic combinations constitute 2.5% of the training set.

#### 2.1.2 Independent dataset of digenic combinations

In order to assess the performance of diVas, we manually curated a high-confidence dataset that encompasses digenic combinations representative of all the known digenic disorders. To be included in the manually curated dataset, digenic combinations should not belong to the diVas’ positive training set, regardless of the associated phenotypes. Initially, we harnessed generative AI techniques to extract phenotype descriptions from an extensive corpus of scientific literature^34^. Subsequently, these extracted phenotypes underwent a rigorous manual curation and validation process led by domain experts. This hybrid approach, combining the efficiency of AI-driven data extraction with the precision of human expertise, resulted in a dataset that captures the complexity of digenic disorders with high confidence. Our dataset comprises over 600 digenic combinations (399 unique genes for a total of 452 unique gene pairs) that were used to assess the accuracy of diVas in terms of pathogenicity and digenic mechanism prediction.

#### 2.1.3 Digenic combinations generation

DiVas has the capability to analyze both Single Nucleotide Variants (SNVs) and Insertion/Deletion (INDEL) variants in both the GRCh37 and GRCh38 genome assemblies.

Each instance created by diVas to represent a digenic combination comprises a minimum of 2 variants, which can be in a homozygous or heterozygous state across both genes. The maximum number of variants in an instance is 4, which occurs in cases of compound heterozygosity on both genes.

In instances of compound heterozygous combinations, the presence of at least one coding variant in each gene is mandatory for inclusion in the analysis. If familial genetic data is accessible, diVas employs this information to effectively reduce the number of instances requiring evaluation. This reduction is achieved by eliminating combinations that do not adhere to the expected pattern of transmission, utilizing genetic information from both affected and unaffected family members. The assumption is that a candidate digenic combination must be shared among affected family members within a family and be absent as a whole from healthy relatives.

In situations where multiple instances have been generated for a particular gene pair, diVas retains at most 4 combinations for each gene pair: for each gene, two inheritance hypotheses are considered (dominant vs recessive) and the variant(s) with the highest pathogenic impact based on the pathogenicity score outlined in section 2.1.4.1 are chosen.

#### 2.1.4 Features set

The feature set employed to characterize each digenic combination is designed to encompass the effects of variants on gene functions, interactions between genes, and associations between genes and phenotypes. The detailed description of the features and the model is reported in Limongelli et al.^35^ Features at variant and gene level describe each gene of the digenic combination independently. Unlike ORVAL and DiGePred, diVas has deliberately moved away from sorting the two genes within a pair based on a single feature or in a random manner. Instead, duplicated variant and gene-level features have been independently arranged according to their minimum and maximum values.

##### 2.1.4.1 Variant impact on gene

The influence of the variant(s) on each gene within the gene pair is quantified using a pathogenicity score^36^. This score incorporates various pieces of information, including the variant’s prevalence in the broader population, *in silico* predictions of pathogenicity, and the classification of the variant in publicly available databases. Specifically, the pathogenicity score assigned to each variant in a digenic pair corresponds to the count of ACMG/AMP criteria that the variant satisfies, as detailed in our earlier work^6,15,36^.

##### 2.1.4.2 Gene-Gene interaction

Several gene-gene level attributes have been incorporated to capture indications of interactions between genes. These attributes encompass biological distance, similarity in pathways based on Gene Ontology, phenotypic similarity, and coexpression. The specific resources and versions utilized are documented in Table 1.

**Table 1:**
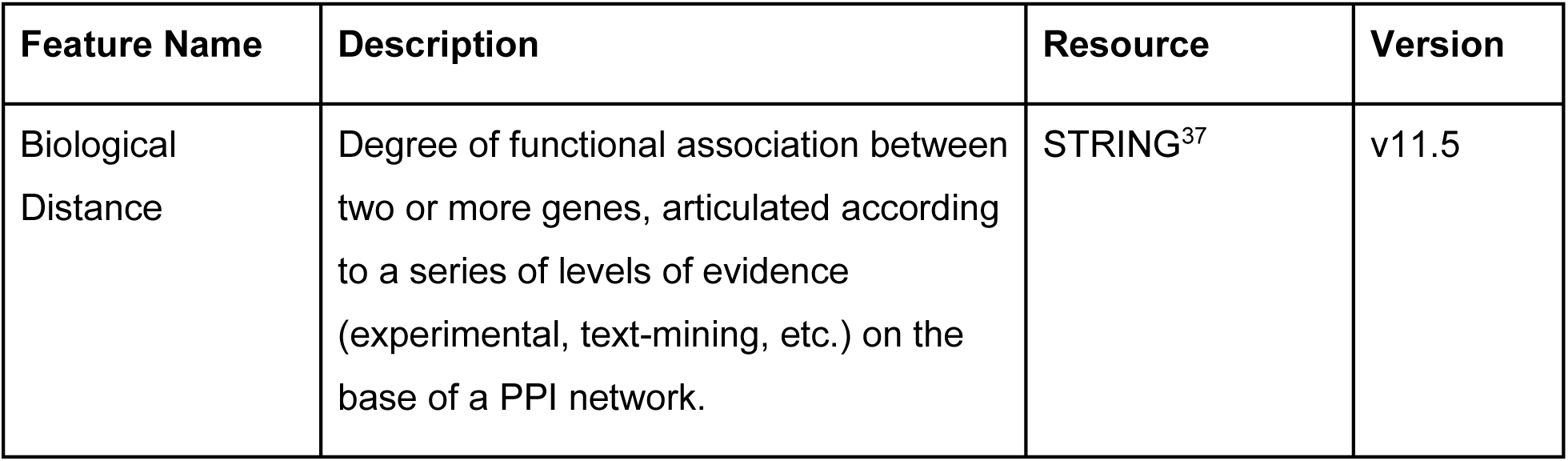
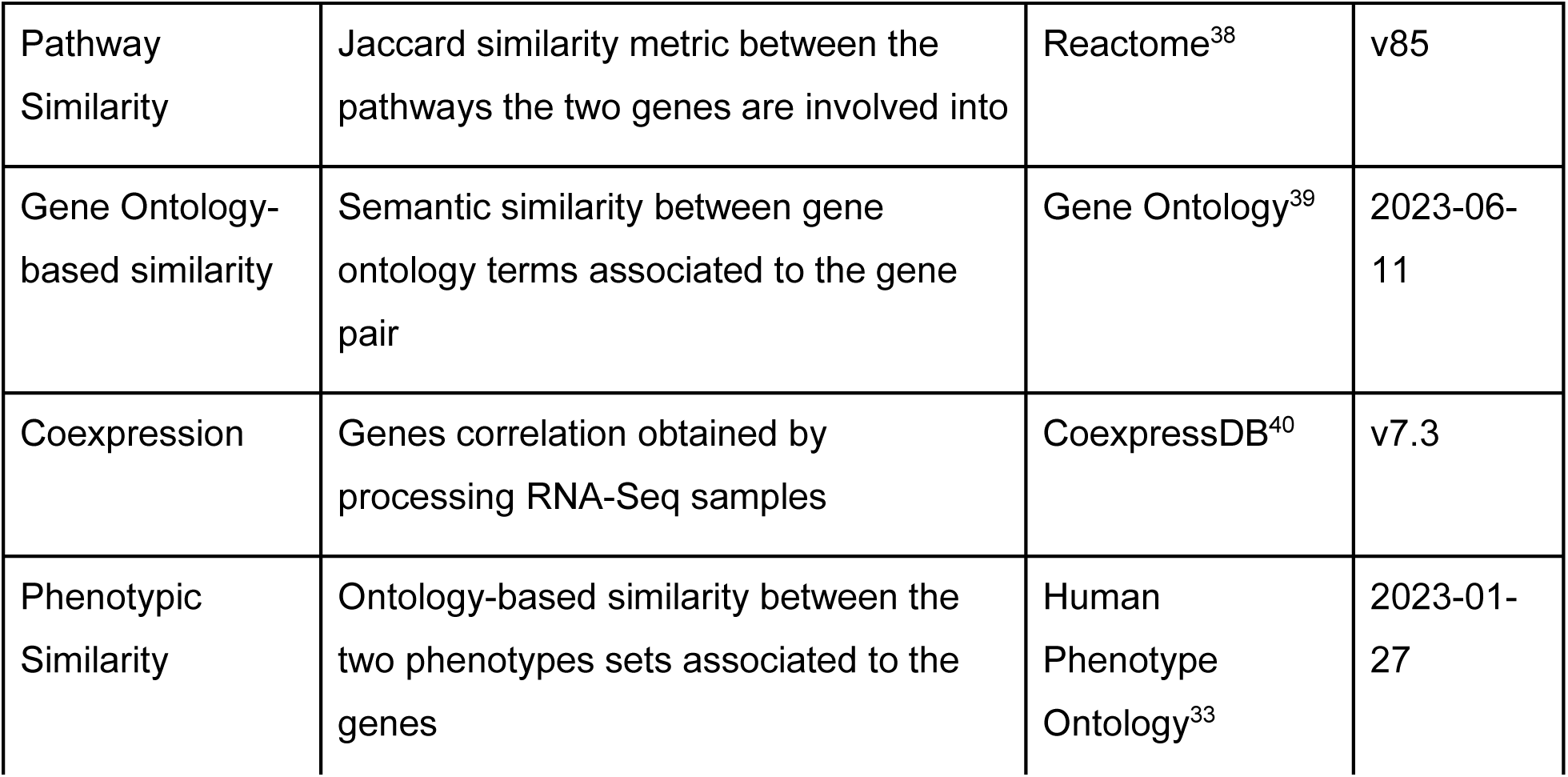
Gene-Gene interaction features. For each attribute, the name, description, associated resource, and version are provided.

| Feature Name | Description | Resource | Version |
| --- | --- | --- | --- |
| Biological Distance | Degree of functional association between two or more genes, articulated according to a series of levels of evidence (experimental, text-mining, etc.) on the base of a PPI network. | STRING<sup>37</sup> | v11.5 |
| Pathway Similarity | Jaccard similarity metric between the pathways the two genes are involved into | Reactome<sup>38</sup> | v85 |
| Gene Ontology-based similarity | Semantic similarity between gene ontology terms associated to the gene pair | Gene Ontology<sup>39</sup> | 2023-06-11 |
| Coexpression | Genes correlation obtained by processing RNA-Seq samples | CoexpressDB<sup>40</sup> | v7.3 |
| Phenotypic Similarity | Ontology-based similarity between the two phenotypes sets associated to the genes | Human Phenotype Ontology<sup>33</sup> | 2023-01-27 |

##### 2.1.4.3 Phenotypic associations

The connection between the gene pair and patient phenotypes described through HPO terms is quantified using diverse metrics. The objective is to prioritize digenic combinations that collectively provide a more robust explanation of the patient’s traits. By computing a measure of similarity between a patient’s clinical manifestations (provided as a set of HPO terms) and disease descriptions associated with genes, phenotype-based prioritization tools utilize standards and quantitative similarity measures to cluster and compare phenotype sets.

The phenotypic similarity score is based on HPO information. It is computed starting from standard phenotypic similarity indexes (the HPOSim package^41^) and leveraging the Resnik similarity between two phenotypic terms and the Best Match Average approach to compute the whole similarity between 2 sets of HPO terms, as described in Deng et al., 2015^41^.

We calculate two distinct phenotypic similarity scores that measure the phenotype similarities between the input set of HPO terms and the HPO terms associated with each gene in the combination. These scores indicate the extent to which each gene in the combination explains the observed phenotypes in the patient.

### 2.2 Classification system

We designed a structured pipeline implementing the diVas algorithm that accepts a list of variants from the proband as input. These variants must be provided in a Variant Call Format (VCF) file, compatible with both the GRCh37 and GRCh38 assemblies. This pipeline performs variant annotation, generates all potential candidate digenic pairs as previously described, and subsequently categorizes each pair as either pathogenic or non-pathogenic, relying on the probability of pathogenicity predicted by the ML model. Our ML classifier, tailored to a “phenotype-driven” approach, is trained using the comprehensive variant-level, gene-level, and phenotype-level feature set detailed in section 2.1.4.

Consequently, the phenotype-driven model requires an additional input: the list of the phenotypes observed in the patient, as HPO terms. However, certain scenarios, such as prenatal testing, may pose challenges in accessing the patient’s phenotypic information. To address such cases, we also introduced a “phenotype-free” model, which relies solely on variant-level and gene-level features, excluding phenotypic information from its input. In the case of the phenotype-driven model, following the prediction process, if a pair is identified as pathogenic, a Bayesian approach that harnesses XAI through SHAP^42,43^ is employed. This approach serves to determine whether the positively predicted pair adheres to a DM mechanism or not (No-DM)^26^ thereby predicting the digenic mechanism. The classification workflow for the phenotype-driven approach is visualized in Figure 2. Further details concerning the ML classifier for digenic prediction and the prediction of the digenic mechanism are elaborated upon in Sections 2.2.1 and 2.2.2.

**Figure 2.**
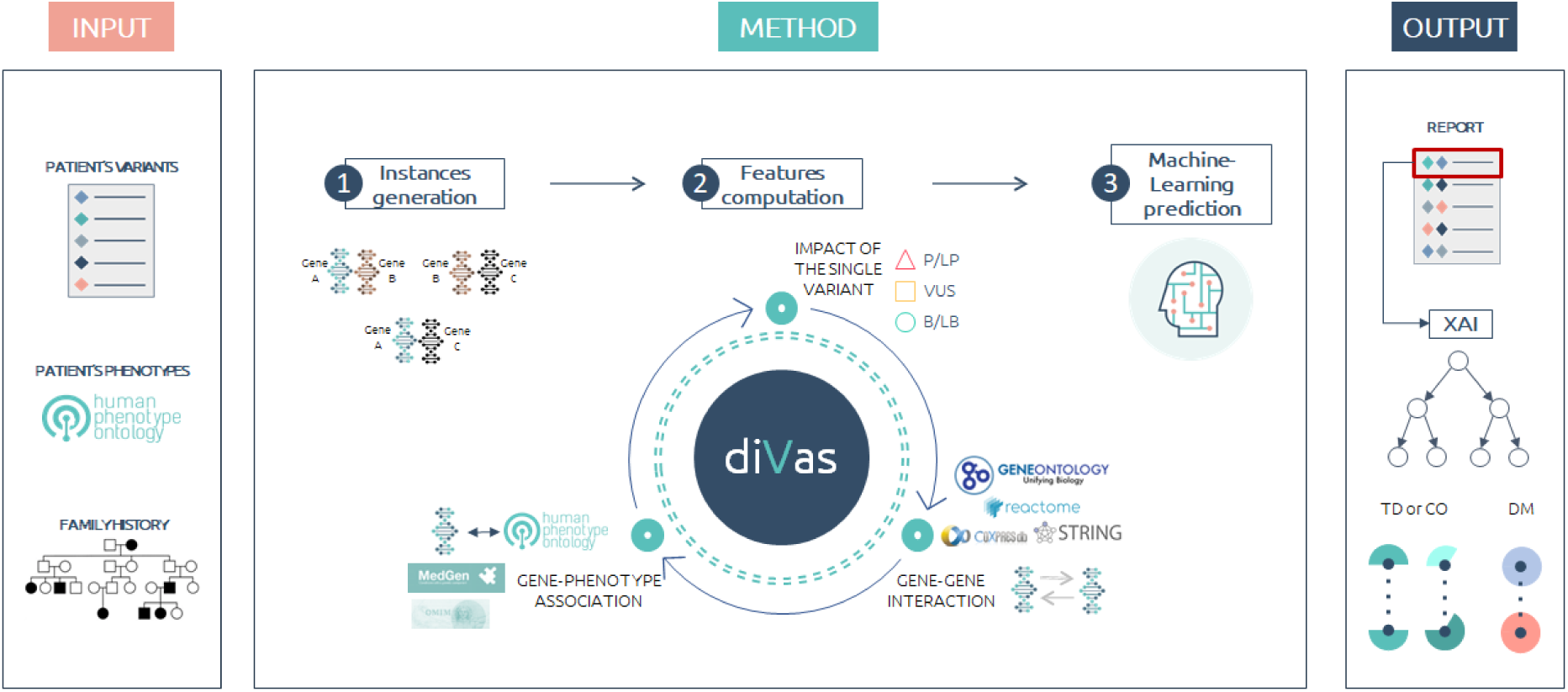
The diVas Workflow Overview: diVas initiates its process by accepting a VCF file, which encompasses the proband’s SNVs and INDELs, along with the patient’s HPO terms and optional family data. This data is then streamlined to curate a list of potential candidate combinations. For every unique instance, a diverse range of features is calculated. Leveraging a pre-trained ML model, each pair is classified as either pathogenic or non-pathogenic, accompanied by a probability score. These combinations are then ranked based on their predicted probabilities. Notably, for every combination identified as pathogenic, diVas conducts an analysis of the ML classifier’s decision pattern, pinpointing the digenic mechanism (through the Bayesian XAI-based digenic effect predictor module) and differentiating between DM combinations and No-DM (both CO and TD) combinations.

#### 2.2.1 Digenic prediction model

We trained the ML model for digenic variant interpretation using a strategy of 500 repeated hold-outs. In each iteration, the dataset was partitioned into training, validation, and test subsets. To minimize potential circularity and avoid overly optimistic outcomes in the test set, we ensured that, during each hold-out, gene pairs in the test subset did not overlap with those in the training or validation subsets^44^. In the training set, we conducted inner cross-validation to determine the optimal hyper-parameters for the ML model. The validation set was employed to identify the best classification threshold. Given the frequent imbalance in variant interpretation datasets, adjusting this threshold can enhance the model’s performance and utility for the minority class^45^.

Then, the best model, using the optimized threshold, predicts the test set, and various performance metrics are evaluated. This process is reiterated 500 times, with the metrics’ statistics averaged over these repetitions. The ultimate classification framework consists of an ensemble of 500 classifiers. For a new instance, the final predicted class is determined by the majority of classifiers, while the predicted probability is the median of all predicted probabilities. Further details on the repeated hold-out procedure are elaborated in the Supplementary Methods (refer to the *DiVas training procedure* under Materials and Methods).

Additionally, to investigate whether the model may have different performance on distinct disorders, we performed a leave-one-phenotype-out validation: we firstly selected a set of different disease categories exploiting Clingen gene-based classification^46^: among all the Clinical Domains curated by Clingen Working Groups, only categories with a minimum of 10 pathogenic combinations belonging to our dataset have been retained (see Table S1). With this approach, we could evaluate 9 out of the 13 available groups from ClinGen. For each disease category, we excluded the pathogenic pairs associated with that category and the negative pairs involving the same genes from the training set. We trained the ML classifier on the remaining set of variants and we evaluated the performance on the excluded pathogenic pairs. With this procedure, we aim at assessing the generalization capability of our approach on instances relative to disorders not represented in the training set.

#### 2.2.2 XAI-based digenic mechanism prediction

As previously mentioned, DI can encompass various disease mechanisms, such as TD, CO, and DM. After a pair is predicted to be pathogenic, understanding its specific subcategory can be valuable.

Distinguishing between a DM and a TD or a CO combination for pathogenic digenic variants is crucial. In a DM, each mutation in different genes is independently pathogenic, leading to two separate genetic conditions. In contrast, in the other digenic mechanisms, two mutations jointly cause a specific phenotype. This distinction is crucial for accurate clinical management, as treatment strategies might differ based on the underlying genetic cause^47^. Moreover, understanding whether it’s a DM or not is essential for genetic counseling, offering insights into inheritance patterns and risks to family members. The differentiation also has significant implications for research, providing valuable information about gene-gene interactions and their role in disease manifestation, and for predicting disease progression and outcomes.

One potential approach is to develop an additional ML classifier, similar to those described in previous studies^21,48^.

Along those lines, authors of previous work identified a set of informative features that can differentiate between DM, TD, and CO subcategories. However, rather than constructing an entirely new ML model based on these distinct features, we adopt a different strategy. We begin by recognizing that positive pairs falling into different subcategories should exhibit varying characteristics. For instance, in the case of DM, we expect that both variants within a pair will possess high pathogenicity scores, while the gene-gene interaction might be minimal. Conversely, in scenarios of TD or CO, we expect strong interconnections between the involved genes. Thus, we propose that the decision-making process of the classifier should diverge for DM and No-DM (TD and CO) scenarios.

XAI is an emerging topic in AI, driven by the need for more trustworthy and transparent classification systems^42^. The aim of XAI is to develop methodologies that support humans to understand the classification process made by complex black-box algorithms, such as deep networks or ensemble methods. Local XAI focuses on providing an explanation for each single prediction made by a ML classifier^49^. We assume that the local explanation of a digenic prediction can uncover the digenic mechanism (DM or No-DM). For example, in the case of a TD pair, we anticipate that the classifier would identify it as positive due to the pronounced gene-gene interaction and phenotypic similarity. On the other hand, for a DM scenario, the decision-making process would likely place greater emphasis on the values of variant-level features. In particular, we used SHAP, a widely applied local XAI method using a game theory approach to dissect the predicted probability and assign a Shapley value to each feature. This numeric value can be interpreted as the contribution of that feature to the predicted probability^50^. To develop our XAI-based digenic effect predictor, we compute the Shapley values for the prediction of each sample with a known digenic effect. Since our final prediction pipeline is an ensemble of 500 classifiers, for a single instance we computed the local XAI of the classifier in the ensemble whose predicted probability for that instance is equal to the median predicted probability. Subsequently, we sum the computed Shapley values for each of the three features category: gene-level features, variant-level features and phenotype-level features. On the training set, we calculate the first percentile, the median and the third percentile of the Shapley values for each group. For each instance, and for each features group, we assign a categorical attribute “low”, “low-median”, “median-high” or “high” if the sum of the Shapley values for that group is below the first percentile, between the first percentile and the median, between the median and the third percentile, or above the third percentile. Therefore, each positive predicted instance will be further characterized by three categorical features: *gene_level_contribution*, *variant_level_contribution* and *pheno_level*_*contribution*, each having 4 different possible values (*low*, *low-median*, *median-high*, *high*). We calculated the probabilities of being DM given each feature contribution to classification, and we used a Naive Bayes approach to classify a new positive predicted instance as follows:

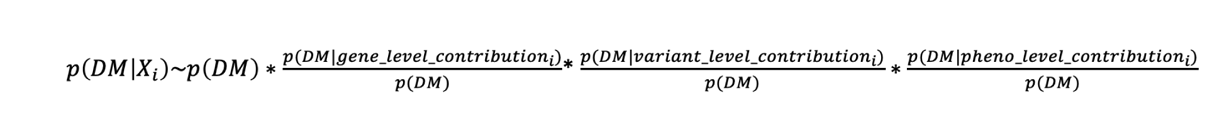

**Equation 1** Naive Bayes predicted probability formula.

Where P(DM) is calculated as the frequency of DM variants in training (28%). The validation strategy on the 500 repeated sampling is described in the Supplementary materials (*XAI-based digenic mechanism prediction: validation approach on training* in supplementary Materials and Methods).

### 2.3 Benchmarking with existing digenic interpretation tools

DiVas performances were compared with other recently developed digenic interpretation tools, namely ORVAL, DiGePred, DIEP, and OligoPVP. While ORVAL, DiGePred, and DIEP were examined in terms of both sensitivity and prioritization capability, OligoPVP’s evaluation focused solely on its prioritization performance due to its lack of binary classification.

The analysis was conducted using data from 11 real cases (gene panel and WES sequencing). These data were collected from participants who provided informed consent for the use of their data for research purposes. The consent process was conducted by the respective partner institutions (see Acknowledgements).

VCF files were pre-filtered to exclude variants with low quality and limited impact on genes. ORVAL integrates the pathogenicity predictor VarCoPP2.0^20^ and has been run from the open platform (https://orval.ibsquare.be/, accessed in August 2023) with default suggested parameters: it filtered out common variants with a MAF>0.035, all intronic variants that have a distance from the exon edge greater than 13 nucleotides and all synonymous variants that have a distance greater than 195 nucleotides from each exon edge. We submitted to ORVAL a tab-delimited file with variant coordinates and associated zygosity.

DiGePred has been executed on a local machine (8 CPU and 16GB RAM) from precomputed scores (download in August 2023) on all human gene pairs available at https://github.com/souhridm/DiGePred. As DiGePred does not take into account single variant information, we provided the list of unique gene pairs for each sample and tested the performance with the default suggested DiGePred model (unaffected_no_gene_overlap), with threshold for classification equal to 0.496, as reported in the paper^23^.

OligoPVP has been run on Amazon Web Service (AWS) cloud EC2 instance (8 CPU and 32GB RAM) based on PhenomeNet Variant Predictor (PVP) v2.1. VCF file with filtered variants and the list of HPO terms describing each patient were provided as input to the tool.

DIEP was run locally (8 CPU and 16GB RAM) using the implementation provided by the authors (https://github.com/pmglab/DIEP). The threshold for classification is 0.5, as reported in the paper^24^.

DiVas was run on AWS cloud EC2 instances based cluster (1-2 CPU and 4-16GB RAM) with family analysis when available, while in single-proband modality for the remaining samples. Benchmark analysis was performed both with the phenotype-driven algorithm and with the phenotype-free model: as the aim of the phenotype-free model is to prioritize digenic pairs that show high-level gene-gene interaction, it has been trained, tested and validated only on samples with a No-DM diagnosis.

Table 2 reports a summary of benchmark tools specifications.

**Table 2:** Release information and availability of tools used for benchmarking.

| Tool | Version | Last release date | WebApp available | Code |
| --- | --- | --- | --- | --- |
| ORVAL/VarCoPP | 3.0.0 | 20-06-2023 | Yes | https://github.com/sofiapapad90/VarCoPP |
| DiGePred | - | 22-09-2021 | Yes | https://github.com/souhridm/DiGePred |
| DIEP | - | 09-08-2022 | No | https://github.com/pmglab/DIEP |
| OligoPVP | 2.1.1 | 14-06-2020 | No | https://github.com/bio-ontology-research-group/phenomenet-vp |
| diVas | 3.0.0 | 01-09-2023 | No | Full description of the method is available.<br>Predictions are available at<br>oliver.engenome.com |

Further comparison analysis was performed between diVas and ORVAL in terms of digenic mechanism prediction.

## 3. Results

### 3.2 Digenic interpretation

#### 3.2.1 Pathogenicity classification

Before analyzing the digenic dataset with digenic-specific variant interpreters, we investigated whether monogenic variant interpretation guidelines alone could support the digenic analysis. We interpreted all single variants in pathogenic pairs according to ACMG/AMP standard guidelines implemented as in our previous study^36^. Therefore, each variant is categorized in the 5 tier system defined by the ACMG/AMP guidelines (see *Monogenic Variant Interpretation* in Supplementary materials). After interpreting each causative training variant according to the ACMG/AMP guidelines, we found that 59% of variants in DM pairs, 47% in CO and 34% in TD are interpreted as Pathogenic/Likely pathogenic, and up to 16% of variants (in case of TD pairs) are Benign/Likely benign (Figure S1). These results confirm the need for the development of digenic-specific guidelines and approaches for interpretation.

On a 500 repeated hold-out validation, the final best performing model selected for digenic pathogenicity prediction is the phenotype-driven Random Forest.

Table 3 presents the mean, median, and percentiles of various pertinent metrics, calculated from the 500 iterations of hold-out validation. DiVas demonstrates commendable performance in detecting both pathogenic (recall) and benign (specificity) cases. When performance is examined in relation to CO, TD, and DM, the average recall for TD is slightly lower (0.864), compared to 0.965 for CO and 0.91 for DM.

**Table 3:** Performance of the diVas algorithm on 500 repeated sampling iterations. The assessment included stratification of recall based on different categories: True Digenic (TD), Dual Molecular Diagnosis (DM), and Composite (CO).

|  | Mean | Median | 25th Percentile | 75th Percentile |
| --- | --- | --- | --- | --- |
| **Specificity** | 0.999 | 0.999 | 0.998 | 1.000 |
| **Recall** | 0.928 | 0.934 | 0.903 | 0.963 |
| **PRC** | 0.979 | 0.982 | 0.969 | 0.992 |
| **Precision** | 0.952 | 0.959 | 0.934 | 0.979 |
| **F1_score** | 0.939 | 0.942 | 0.923 | 0.959 |
| **Balanced Accuracy** | 0.963 | 0.966 | 0.951 | 0.981 |
| **TD recall** | 0.864 | 0.895 | 0.813 | 0.962 |
| **DM recall** | 0.910 | 0.917 | 0.857 | 1.000 |
| **CO recall** | 0.965 | 1.000 | 0.933 | 1.000 |

Notably, when stratified across digenic mechanisms, the phenotype-driven model exhibits superior performance compared to the phenotype-free model (see Table S2). The DM mechanism was not considered for the phenotype-free model. Specifically, the average recall for TD is 0.864 with the phenotype-driven approach, compared to 0.731 with the phenotype-free approach. On the other hand, the phenotype-driven approach achieves an average sensitivity of 0.965 for CO, which decreases to 0.899 with the phenotype-free approach. The distribution of metrics across the 500 sampling iterations can be observed in Figure S2 and Figure S3.

The model’s generalization capabilities were assessed using the “leave-one-phenotype-out” approach. Table S1 shows the number of True Positive and False Negative pairs in 9 different disorder categories. DiVas shows high recall for the disease category, in particular in Kidney Disease, Inborn Errors of Metabolism and Ocular disorders. The lower recall (0.917) is shown on Skeletal Disorders. Overall, these results suggest that diVas classification is reliable even when the prediction is made on variant pairs associated with disorders not represented in the training dataset. Having a good generalization ability for populations not represented in the training data is crucial for high-stakes applications like healthcare^51^.

#### 3.2.2 Digenic mechanism prediction with XAI

Table S3 reports the Naive Bayes conditional probabilities computed on the positive training set. For each features subgroups (*gene-level, variant-level and phenotype-leve*l), the categorical value (*low, low-median, median-high, high)*, representing the contribution of that features subgroup to classification (proportional to the SHAP values), is reported, along with the conditional probability for DM and No-DM classification. We observe that there’s a 94% probability of a pair being classified as DM when there’s a low contribution from the gene-level feature. Since gene level features broadly represent the gene-gene interactions, this result is in line with what we would expect for DM pairs, in which two pathogenic variants for two independent disorders are simultaneously detected in proband’s genome and no interaction is expected *a priori* between the impacted genes. On the contrary, the probability of being DM given a low variant-level contribution is low. Also in this case the XAI classification pattern follows what we would expect: variants in DM should exhibit high pathogenicity, whereas TD and CO may consist of variants that are less pathogenic when considering a monogenic hypothesis with genes that have a strong interaction (Figure S1). Therefore, the variant-level contribution to TD classification is expected to be low.

On the 500 repeated hold-out validation, performances are reported in Figure 3. In median, the percentage of DM variants in the test folds is 28%, in line with the general proportion of DM in the complete set.

**Figure 3:**
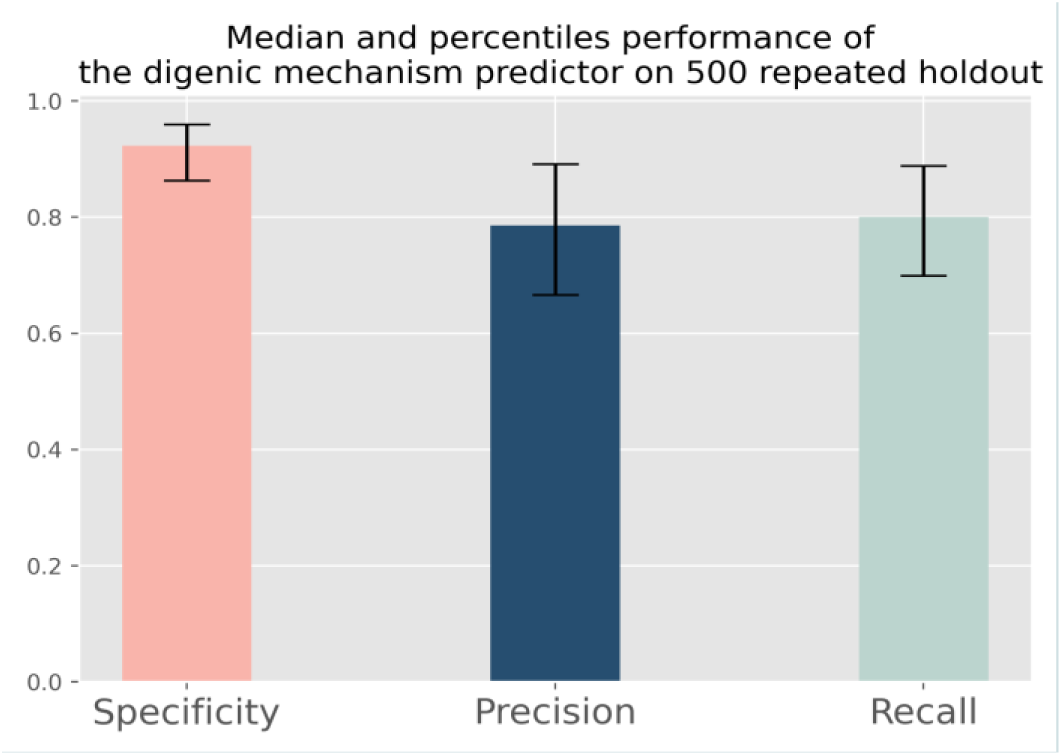
Median, 25th and 75th percentile of specificity, precision and recall of the digenic mechanism predictor on 500 repeated holdout validation. Positive class is DM (Dual Molecular).

### 3.3 Validation on real cases and benchmark analysis

DiVas has been validated on an independent dataset comprising 11 gene panel/WES samples with confirmed digenic diagnoses collected through international collaboration (see *Acknowledgements*). Samples included in the validation set present heterogeneous phenotypes, ranging from skeletal to kidney disorders, hearing impairment, ocular, and cardiology abnormalities. The number of HPO terms describing patients’ traits varies from a minimum of 1 to a maximum of 4 terms (Table S4), with a median of 2. A total of 6 samples have been analyzed in single-proband modality, while for the remaining samples family information has been integrated. For 10 out of 11 pathogenic digenic combinations the digenic mechanism was known *a priori* (4 DM and 6 TD/CO).

Since diVas, ORVAL, OligoPVP and DiGePred adopt different filtering and pair generation strategies, the number of digenic instances evaluated by each tool is different. As a matter of fact, ORVAL predicts all possible variant combinations, while diVas evaluates, for a given gene pair, at most 4 combinations with the more pathogenic variants in those genes (according to different inheritance hypotheses). On the contrary, DiGePred and DIEP do not consider variant pairs and predict the pathogenicity at the gene pair level. Finally, OligoPVP predicts the pathogenicity of each individual variant, subsequently combining variants on genes with evidence of interaction, thus potentially evaluating more than one instance per gene pair. Therefore, the number of combinations ranked by diVas, DiGePred and DIEP is lower potentially enhancing their prioritization ability when compared with ORVAL. For this reason, we evaluated ORVAL prioritization both in default mode and by de-duplicating gene pairs. In this case, for each gene pair, the variant combination predicted with the highest probability by ORVAL is selected. Summary about tools’ performances on the real samples are reported in Table 4 and Figure 4, while Table S5 contains single samples’ detailed statistics. Table S5 also compares diVas with its phenotype-free version to evaluate the impact of a patient’s phenotypes on classification.

**Figure 4.**
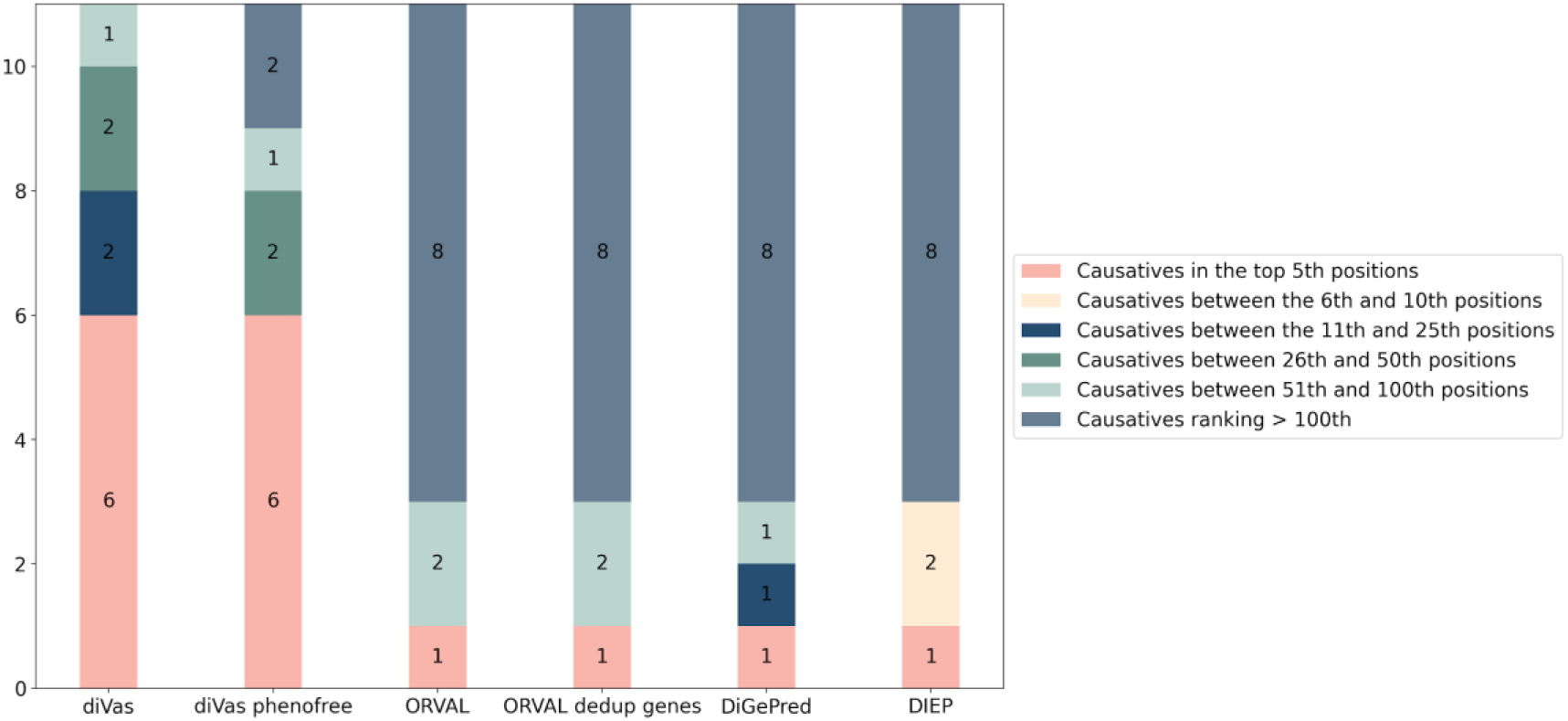
Performance of different tools on digenic pairs prioritization of real cases. For each tool, we report the number of real cases for which the causative pair is ranked 1) in the top 5th positions, 2) between the 6th and 10th 3) between the 11th and 25th, 4) between the 26th and 50th 5) between the 51th and 100th or 6) above the 100th position.

**Table 4:** Median performance of digenic prediction tools on the validation set of real cases. Median ranking of the causative digenic combination, median number of predicted pathogenic and of evaluated digenic combinations is reported. Additionally, we report the sensitivity, computed as the number of true pathogenic pairs correctly classified by the tool divided by the total number of pathogenic cases (11).

| **Tool** | **Median Ranking** | **Sensitivity (fraction of pathogenic pairs correctly classified)** | **Median Number of Predicted Pathogenic combinations ( % over total number of evaluated instances)** | **Median number of evaluated digenic instances** |
| --- | --- | --- | --- | --- |
| **diVas** | 3 | 0.73 | 16 (0.2%) | 6656 |
| **ORVAL** | 154.5 | 0.64 | 3510.5(26,15%) | 13427 |
| **ORVAL_dedup_genes** | 154.5 | 0.64 | 3460(26.33%) | 13143 |
| **DiGePred** | 760 | 0.18 | 30 (0.5%) | 5856 |
| **DIEP** | 346 | 0.55 | 306 (5.23%) | 5855 |
| **diVas phenotype-free** | 5 | 0.73 | 158 (2.37%) | 6656 |

Overall, diVas shows better performance than the other tools in terms of sensitivity, median false positive rate, and median ranking of the causative digenic combination. ORVAL has a sensitivity similar to diVas. However, it predicts nearly 30% of the evaluated instances as pathogenic, failing to adequately control the number of false positives, which is essential for a method used in realistic clinical settings. Also DIEP shows a high number of false positives (Table S5). Deduplicating ORVAL results by gene pair slightly improves its prioritization statistics. While DiGePred demonstrates greater specificity than ORVAL, it exhibits the lowest sensitivity, with only 2 of the 11 causative digenic combinations accurately predicted as pathogenic.

OligoPVP identifies only 2 out of 11 causative combinations, with a median of 102 prioritized pathogenic combinations identified per sample. This outcome might be attributed to the outdated resources used to assess gene-gene interactions. A deeper investigation of the causes of its high missing rate highlights that only 2 out of 11 causative gene pairs are considered as interacting by OligoPVP, and thus used to build the final results. Hopefully, updated gene-gene interaction resources may dramatically impact on OligoPVP’s prioritization performances. For this reason, we decided to exclude OligoPVP from the benchmark analysis reported in Table 4 and Figure 4.

In addition, we compared diVas and ORVAL in terms of digenic sub-classification ability. The diVas XAI-based digenic mechanism predictor correctly subclassified 6 out of 8 digenic combinations that have been previously classified as pathogenic by the tool, resulting in a sensitivity of 75%. ORVAL, aggregating TD/CO in order to be comparable with our classification, correctly subclassified 3 out of 6 predicted pathogenic combinations (Table S6).

### 3.4 Classification performance on selected cases from OLIDA database

We additionally benchmarked diVas on the manually curated high-confidence database described in section 2.1.2. DiVas achieves a sensitivity of 0.81 when applied to 645 causative digenic combinations using the phenotype-driven approach, while it reaches a lower sensitivity of 0.61 when employing the phenotype-free approach.

The XAI-driven digenic mechanism predictor reaches an accuracy of 76% in distinguishing DM and TD/CO, when applied to a subset of 46 causative combinations for which the digenic mechanism was known.

### 3.5 OliVer: sharing a Dataset of Pathogenic Digenic Combinations

We’ve made the manually curated dataset publicly available via a dedicated web app accessible at oliver.engenome.com.

We’ve predicted 645 instances (not used for training purposes) and have made the prediction outcomes available via the OliVer platform (oliver.engenome.com). For every combination, we provide coordinates of manually curated variants, a list of associated phenotypes we’ve curated, and prediction insights from diVas and XAI-driven digenic mechanism predictor. A platform overview is shown in Fig. 5.

**Figure 5:**
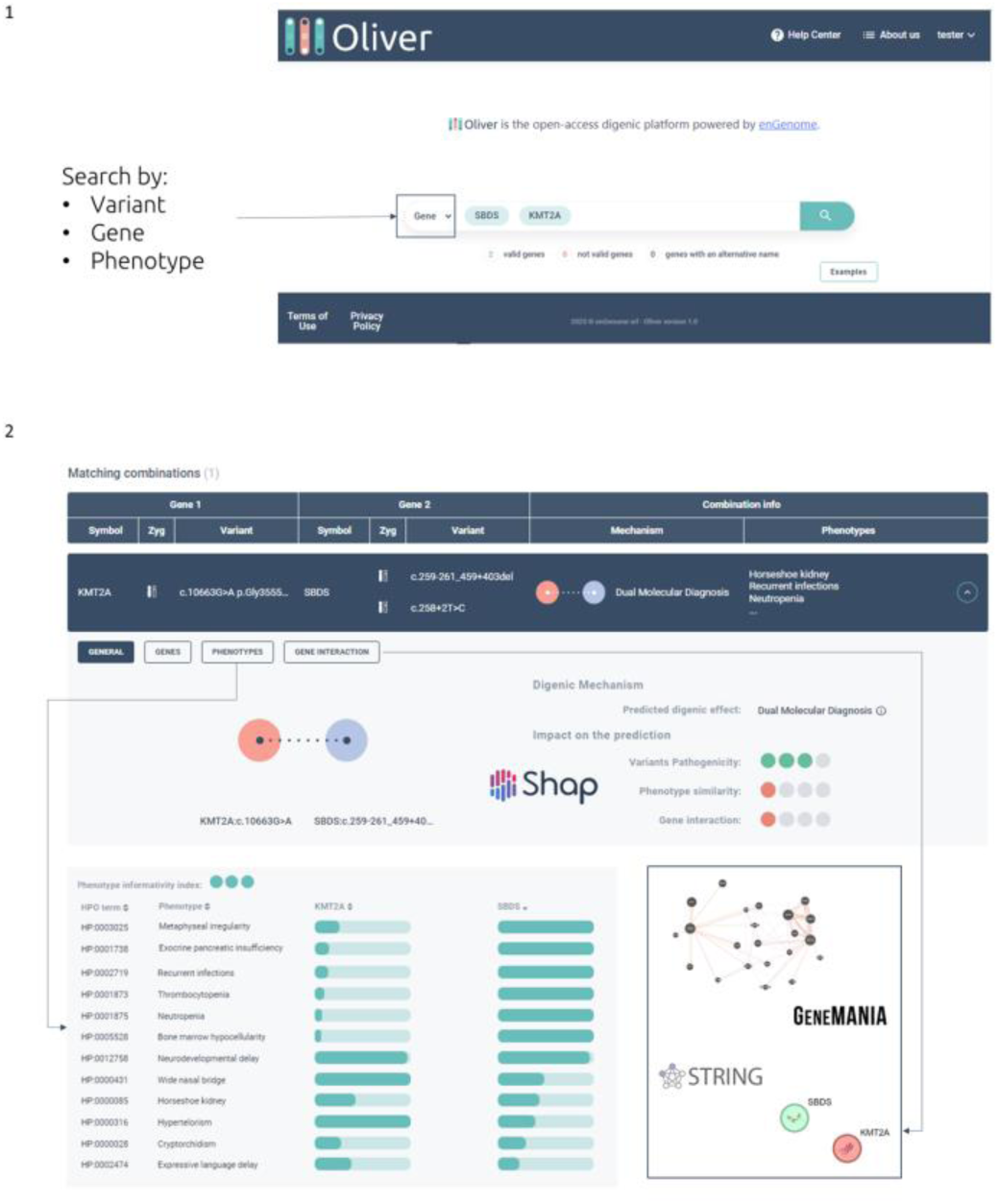
preview of the OliVer platform available at oliver.engneome.com. Panel 1: search tab (by variant, by gene or by phenotype). Panel 2: example of variant search and insight of a combination.

## 4. Discussion

Currently, the diagnostic yield of RDs is on average far below 50%^1^ and there is an urgent need to uplift this value and end the diagnostic odyssey for thousands of patients. One of the open challenges in the field of genomics is to go beyond the “one gene, one disease” paradigm that led the diagnostic genetic approach in the last 50 years^52^. Oligogenic inheritance has gained increasing attention within the ongoing challenge of identifying the causes of rare genetic disorders.

While bioinformatics and computational approaches have greatly simplified and supported the diagnostic process, still the majority of available tools are applied to each single variant in a genome^13,15–18^, without considering interactions between different elements, such as genes and their products. In the oligogenic hypothesis, the genetic cause of a disorder is attributed to the interaction of different mutated genes. As a consequence, currently available approaches for variant interpretation are not suitable for the identification of oligogenic causative variants. The development of methodologies for the interpretation of oligogenic disorders is hampered by limited availability of bona fide pathogenic oligogenic combinations. Unlike ClinVar^28^, a public repository that currently gathers thousands of interpretations under the monogenic hypothesis, the current available database of pathogenic oligogenic hypothesis (OLIDA) gathers around 1600 instances, the majority of them represented by digenic combinations^27^. Another fundamental aspect is that OLIDA is a manually-curated database that gathers oligogenic pathogenic instances from the literature, while ClinVar allows for the submission of clinically-relevant interpretations from submitters worldwide.

Early methods focusing on digenic variant interpretation (like Orval, DiGePred, OligoPVP and DIEP) were all trained on DIDA, the previous version of OLIDA, gathering around 250 pathogenic combinations^22,23^. Orval recently integrated the positive training set of the ML model with the most recently released Olida data. Due to limited availability of bona-fide pathogenic combinations, external validation of such tools can be hampered by circularity issues^44^. The performed benchmark analysis showed that no single tool emerges as a universal standard: while Orval exhibits high sensitivity, its specificity is notably compromised. In contrast, DigePred and DIEP provide greater specificity but are constrained by their gene-level predictions, without considering the variants included in each combination. OligoPVP, currently, is the only published tool that incorporates patient phenotypes into its predictive framework; however, its sensitivity is hindered by outdated gene-gene interaction resources.

Here, we present diVas, an ML-based approach for digenic variant interpretation aiming to overcome the limitations of the other tools described above. Unlike other tools, diVas leverages proband’s phenotypic information to predict the probability of each pair to be causative. DiVas was trained on more than 350 bona fide curated digenic instances. It exploits variant pathogenicity, computed according to standard variant interpretation guidelines through the eVai software^36^ together with gene-gene interaction and gene-phenotype association information, to predict the pathogenicity probability. This value is exploited to rank all the combinations, potentially thousands in a WGS/WES experiment. In 11 clinical cases, we demonstrated that diVas outperformed existing methods in prioritizing the causative pairs in the top positions. The causative pair median ranking is 3 for diVas, with at least a 50-fold reduction if compared with other existing solutions. Better performances have been noticed when a patient’s phenotypic description is provided to the algorithm and the tool proved to generalize well across heterogeneous phenotypic spectrum of different diseases; however, even when the phenotypic description is missing, the phenotype-free model can help in the prioritization of the causative digenic combination. Additionally, diVas is able to account for family segregation, an essential information for clinical utility^53^. An additional layer of explainability (XAI) allows our tool to further discern the digenic disease mechanisms. The fact that our ML model is explainable is an important aspect to promote trust in Artificial Intelligence, and it is highly encouraged by recent EU guidelines on AI^54^. A limitation of our current methodology to predict digenic mechanisms is its resolution: unlike previous work, our subclassification system is binary, thus predicting DM or No-DM cases. A future development will be a subclassification system able to distinguish between CO and TD.

The natural extension of diVas is to move towards oligogenic interpretation. In this case, the challenge is to gain knowledge on the molecular interactions among sets of genes and to obtain a sufficiently robust ground-truth dataset to benchmark and validate such a solution.

The dramatic improvement in prioritization performances achieved by diVas opens the way to the wide-spread usage of these tools on undiagnosed patients. It is possible to imagine a near future where diVas may be exploited in clinical settings on top of routinely adopted diagnostic solutions based on a monogenic approach on deep-phenotyped patients lacking a molecular diagnosis.

In this context, digenic inheritance understanding is becoming crucial. We’ve released a curated dataset of digenic combinations and associated predictions on the OliVer platform (oliver.engenome.com) to advance this understanding. This dissemination aims to highlight digenic inheritance complexities, thereby deepening the scientific community’s foundational knowledge.

Furthermore, providing our curated data and prediction results supports researchers in their endeavors, reflecting our commitment to advancing the field. This resource is a powerful tool for hypothesis generation, findings validation, and new investigations. Beyond data sharing, we aspire to foster a community of researchers and clinicians with a shared interest in digenic inheritance, promoting collaboration, discussion, and insight exchange through the OliVer platform. Our contribution is not just data presentation; it’s an invitation to collaborative scientific exploration, expected to drive further advancements in understanding digenic inheritance and its various implications.

## Availability

oliver.engenome.com

## List of Abbreviations

| Abbreviation | Definition |
| --- | --- |
| 1KGP | 1000 Genomes Project |
| ACMG | American College of Medical Genetics |
| AMP | Association for Molecular Pathology |
| AWS | Amazon Web Services |
| CO | Composite |
| DE | Digenic Effect |
| DI | Digenic Inheritance |
| DM | Dual Molecular Diagnosis |
| INDEL | Insertion/Deletion |
| HPO | Human Phenotype Ontology |
| ML | Machine Learning |
| SNVs | Single Nucleotide Variants |
| TD | True Digenic |
| VCF | Variant Call Format |
| VUS | Variant of Uncertain Significance |
| WES | Whole Exome Sequencing |
| WGS | Whole Genome Sequencing |
| XAI | Explainable Artificial Intelligence |

## Acknowledgements

We would like to extend our gratitude to the following scientists, listed in alphabetical order: Dr. Alfredo Brusco and PhD Lisa Pavinato (University of Turin, Turin, Italy); Dr. Cesare Danesino, Dr. Antonella Minelli, and PhD Ibrahim Taha (University of Pavia, Pavia, Italy); Dr. Edoardo Errichiello and PhD student Mauro Lecca (University of Pavia, Pavia, Italy); Dr. Paolo Gasparini, Dr. Giorgia Girotto, and PhD Anna Morgan (Institute for Maternal and Child Health - IRCCS Burlo Garofolo, Trieste, Italy); Dr. Alessandro Geroldi (University of Genoa, Genoa, Italy); Dr. Corrado Mammì and Dr. Manuela Priolo (Hospital of Reggio Calabria Az. Ospedaliera Bianchi-Melacrino-Morelli, Reggio Calabria, Italy); Dr. Nikolaos M. Marinakis, Dr. Joanne Traeger-Synodinos, and PhD Faidon-Nikolaos Tilemis (National and Kapodistrian University of Athens, Athens, Greece); Dr. Marco Tartaglia (IRCCS Bambino Gesù Children’s Hospital, Rome, Italy); Dr. Enza Maria Valente and PhD Valentina Serpieri (IRCCS Mondino Foundation, Pavia, Italy).

We also extend our thanks to Anne Bowcock (PhD - Icahn School of Medicine at Mount Sinai) for her invaluable support in data curation.

## References

1. 100,000 Genomes Pilot on Rare-Disease Diagnosis in Health Care — Preliminary Report. *New England Journal of Medicine* 385, 1868–1880 (2021).

2. Marwaha, S., Knowles, J. W. & Ashley, E. A. A guide for the diagnosis of rare and undiagnosed disease: beyond the exome. Genome Medicine 14, 23 (2022).

3. Álvarez-Mora, M. I. et al. Diagnostic yield of next-generation sequencing in 87 families with neurodevelopmental disorders. Orphanet Journal of Rare Diseases 17, 60 (2022).

4. Frésard, L. & Montgomery, S. B. Diagnosing rare diseases after the exome. *Cold Spring Harb Mol Case Stud* 4, a003392 (2018).

5. Vinkšel, M., Writzl, K., Maver, A. & Peterlin, B. Improving diagnostics of rare genetic diseases with NGS approaches. J Community Genet 12, 247–256 (2021).

6. Sarah L. Stenton et al. Critical assessment of variant prioritization methods for rare disease diagnosis within the Rare Genomes Project. medRxiv 2023.08.02.23293212 (2023) doi:10.1101/2023.08.02.23293212.

7. Rahit, K. M. T. H. & Tarailo-Graovac, M. Genetic Modifiers and Rare Mendelian Disease. *Genes (Basel)* **11**, 239 (2020).

8. Richards, S. et al. Standards and guidelines for the interpretation of sequence variants: a joint consensus recommendation of the American College of Medical Genetics and Genomics and the Association for Molecular Pathology. Genet Med 17, 405–424 (2015).

9. Deltas, C. Digenic inheritance and genetic modifiers. *Clin Genet* **93**, 429–438 (2018).

10. König, E. et al. Exploring digenic inheritance in arrhythmogenic cardiomyopathy. BMC Medical Genetics 18, 145 (2017).

11. Schäffer, A. A. Digenic inheritance in medical genetics. J Med Genet 50, 641–652 (2013).

12. Flygare, S. et al. The VAAST Variant Prioritizer (VVP): ultrafast, easy to use whole genome variant prioritization tool. BMC Bioinformatics 19, 1–13 (2018).

13. Jaganathan, K. et al. Predicting Splicing from Primary Sequence with Deep Learning. Cell 176, 535–548.e24 (2019).

14. Limongelli, I., Marini, S. & Bellazzi, R. PaPI: pseudo amino acid composition to score human protein-coding variants. BMC Bioinformatics 16, 123 (2015).

15. Nicora, G., Zucca, S., Limongelli, I., Bellazzi, R. & Magni, P. A machine learning approach based on ACMG/AMP guidelines for genomic variant classification and prioritization. Sci Rep 12, 2517 (2022).

16. Rentzsch, P., Witten, D., Cooper, G. M., Shendure, J. & Kircher, M. CADD: predicting the deleteriousness of variants throughout the human genome. Nucleic Acids Research 47, D886–D894 (2019).

17. Ioannidis, N. M. et al. REVEL: An Ensemble Method for Predicting the Pathogenicity of Rare Missense Variants. Am J Hum Genet 99, 877–885 (2016).

18. Yuan, X. et al. Evaluation of phenotype-driven gene prioritization methods for Mendelian diseases. Brief Bioinform 23, bbac019 (2022).

19. Okazaki, A. & Ott, J. Machine learning approaches to explore digenic inheritance. Trends in Genetics (2022) doi:10.1016/j.tig.2022.04.009.

20. Papadimitriou, S. et al. Predicting disease-causing variant combinations. Proc. Natl. Acad. Sci. U.S.A. 116, 11878–11887 (2019).

21. Gazzo, A. et al. Understanding mutational effects in digenic diseases. Nucleic Acids Research 45, e140–e140 (2017).

22. Renaux, A. et al. ORVAL: a novel platform for the prediction and exploration of disease-causing oligogenic variant combinations. Nucleic Acids Research 47, W93–W98 (2019).

23. Mukherjee, S. et al. Identifying digenic disease genes via machine learning in the Undiagnosed Diseases Network. The American Journal of Human Genetics 108, 1946–1963 (2021).

24. Yuan, Y., Zhang, L., Long, Q., Jiang, H. & Li, M. An accurate prediction model of digenic interaction for estimating pathogenic gene pairs of human diseases. Computational and Structural Biotechnology Journal (2022) doi:10.1016/j.csbj.2022.07.011.

25. Boudellioua, I., Kulmanov, M., Schofield, P. N., Gkoutos, G. V. & Hoehndorf, R. OligoPVP: Phenotype-driven analysis of individual genomic information to prioritize oligogenic disease variants. Sci Rep 8, 14681 (2018).

26. Gazzo, A. M. et al. DIDA: A curated and annotated digenic diseases database. Nucleic Acids Research 44, D900–D907 (2016).

27. Nachtegael, C. et al. Scaling up oligogenic diseases research with OLIDA: the Oligogenic Diseases Database. Database 2022, baac023 (2022).

28. Landrum, M. J. et al. ClinVar: public archive of relationships among sequence variation and human phenotype. Nucleic Acids Res 42, D980–D985 (2014).

29. Yang, R. et al. Case Report: Expanding the Digenic Variants Involved in Thyroid Hormone Synthesis−10 New Cases of Congenital Hypothyroidism and a Literature Review. Front. Genet. 12, 694683 (2021).

30. Iafusco, F. et al. NGS Analysis Revealed Digenic Heterozygous GCK and HNF1A Variants in a Child with Mild Hyperglycemia: A Case Report. Diagnostics 11, 1164 (2021).

31. Chen, Q. et al. Digenic Variants in the TTN and TRAPPC11 Genes Co-segregating With a Limb-Girdle Muscular Dystrophy in a Han Chinese Family. Front. Neurosci. 15, 601757 (2021).

32. Posey, J. E. et al. Resolution of Disease Phenotypes Resulting from Multilocus Genomic Variation. N Engl J Med 376, 21–31 (2017).

33. Köhler, S. et al. The Human Phenotype Ontology in 2021. Nucleic Acids Research 49, D1207–D1217 (2021).

34. Feng, Y., Qi, L. & Tian, W. PhenoBERT: A Combined Deep Learning Method for Automated Recognition of Human Phenotype Ontology. IEEE/ACM Trans Comput Biol Bioinform 20, 1269–1277 (2023).

35. Limongelli, I., et al. Metodo predittivo per determinare la patogenicità di combinazioni di varianti digeniche o oligogeniche. (2022).

36. Nicora, G. et al. CardioVAI: An automatic implementation of ACMG-AMP variant interpretation guidelines in the diagnosis of cardiovascular diseases. Human Mutation 39, 1835–1846 (2018).

37. Szklarczyk, D. et al. STRING v11: protein-protein association networks with increased coverage, supporting functional discovery in genome-wide experimental datasets. Nucleic Acids Res 47, D607–D613 (2019).

38. Gillespie, M. et al. The reactome pathway knowledgebase 2022. Nucleic Acids Res 50, D687–D692 (2022).

39. Carbon, S. & Mungall, C. Gene Ontology Data Archive. (2018) doi:10.5281/ZENODO.2529950.

40. Obayashi, T., Kagaya, Y., Aoki, Y., Tadaka, S. & Kinoshita, K. COXPRESdb v7: a gene coexpression database for 11 animal species supported by 23 coexpression platforms for technical evaluation and evolutionary inference. Nucleic Acids Res 47, D55–D62 (2019).

41. Deng, Y., Gao, L., Wang, B. & Guo, X. HPOSim: an R package for phenotypic similarity measure and enrichment analysis based on the human phenotype ontology. PLoS One 10, e0115692 (2015).

42. Caruana, R., Lundberg, S., Ribeiro, M. T., Nori, H. & Jenkins, S. Intelligible and Explainable Machine Learning: Best Practices and Practical Challenges. in Proceedings of the 26th ACM SIGKDD International Conference on Knowledge Discovery & Data Mining 3511–3512 (Association for Computing Machinery, 2020). doi:10.1145/3394486.3406707.

43. A, S. & R, S. A systematic review of Explainable Artificial Intelligence models and applications: Recent developments and future trends. Decision Analytics Journal 7, 100230 (2023).

44. Grimm, D. G. et al. The evaluation of tools used to predict the impact of missense variants is hindered by two types of circularity. Hum Mutat 36, 513–523 (2015).

45. Zou, Q., Xie, S., Lin, Z., Wu, M. & Ju, Y. Finding the Best Classification Threshold in Imbalanced Classification. Big Data Research 5, 2–8 (2016).

46. Rivera-Muñoz, E. A. et al. ClinGen Variant Curation Expert Panel experiences and standardized processes for disease and gene-level specification of the ACMG/AMP guidelines for sequence variant interpretation. Hum Mutat 39, 1614–1622 (2018).

47. Kadlubowska, M. K. & Schrauwen, I. Methods to Improve Molecular Diagnosis in Genomic Cold Cases in Pediatric Neurology. Genes (Basel) 13, 333 (2022).

48. Versbraegen, N. et al. Using game theory and decision decomposition to effectively discern and characterise bi-locus diseases. Artificial Intelligence in Medicine 99, 101690 (2019).

49. Guidotti, R. et al. A Survey of Methods for Explaining Black Box Models. ACM Comput. Surv. 51, 93:1–93:42 (2018).

50. Lundberg, S. M. et al. From local explanations to global understanding with explainable AI for trees. Nat Mach Intell 2, 56–67 (2020).

51. Kelly, C. J., Karthikesalingam, A., Suleyman, M., Corrado, G. & King, D. Key challenges for delivering clinical impact with artificial intelligence. BMC Med 17, 1–9 (2019).

52. Rimoin, D. L. & Hirschhorn, K. A History of Medical Genetics in Pediatrics. Pediatr Res 56, 150–159 (2004).

53. Taha, I. et al. Phenotypic Variation in Two Siblings Affected with Shwachman-Diamond Syndrome: The Use of Expert Variant Interpreter (eVai) Suggests Clinical Relevance of a Variant in the KMT2A Gene. Genes (Basel) 13, 1314 (2022).

54. The Act. *The Artificial Intelligence Act* https://artificialintelligenceact.eu/the-act/ (2021).

