## Supplementary File 1 for "Digenic variant interpretation with hypothesis-driven explainable AI"

### Supplementary Materials

#### Materials and Methods

##### DiVas training procedure

At each step of the 500 repeated hold-out validation, we select 65% of data for training, 20% for validation and the remaining 15% for test. Each subset is stratified, meaning that the proportion of classes is the same (2.5% of pathogenic) as in the complete dataset. Additionally, to reduce possible circularity and over-optimistic results on the test set, at a given step, the test set has no overlap in terms of gene pairs with instances belonging to training or validation set. On the training set, 5-fold inner cross validation is performed to select the best hyperparameters (i.e. the model that results in the highest F1 score, harmonic mean between precision and recall) for each ML classifier that we tested. Subsequently, the classifier trained with the best hyperparameters is used to predict the validation set. On these instances, we also select the best threshold for classification. Usually, an ML predictor classifies as positive all instances whose predicted probability is greater than 0.5. The 0.5 threshold can work well on balanced problems, when the class distribution is the same. On the contrary, herefore, we select the threshold that maximizes the F1 score, on the validation predictions. Finally, we evaluate the performance of the models with their optimized thresholds on the test set. Both for the phenotype-driven and the phenotype-free approach, we tested different ML classifiers, such as Random Forest, Logistic Regression, and Gradient Boosting. For the diVas phenotype-free approach, the training procedure was the same. In this case, however, we decided to train the models on True Digenic (TD) and Composite (CO) pairs alone: in fact, we observed that in the case of Dual Molecular Diagnosis (DM) the values of the gene-level features are low, since variants in DM pairs occur in genes that are not involved in the same molecular pathways. By also adding DM to the training set, the model would have been biased towards positive classification even when the gene-level features contribution is low.

##### XAI-based digenic mechanism prediction: validation approach on training

To evaluate the effectiveness of the SHAP-based Naive Bayes within the repeated hold-out validation, we applied the following strategies:

1. For each seed, we calculate the SHAP values on the validation set for predicted positive samples with a digenic effect (DM or No-DM) available

2. On the validation set, we calculated the gene_level, variant_level and phenotype_level contribution as the sum of the SHAP values. We then calculated the median and percentile to categorize such contributions as described in the main manuscript. We calculated the probability of being DM and the conditional probabilities on such a set of data.
3. For each test sample predicted as positive and with available digenic effect, we calculated the posterior probability of being DM according to the Naive Bayes approach described in Equation 1. On these predictions, we evaluated different metrics, such as precision in identifying DM pairs, recall and specificity.

### Results

#### diVas performance on 500 repeated hold-out validation

| **Disorder category** | **# combinations** | **RF confusion matrix TP(FN)** | **RF recall** |
|---|---|---|---|
| Kidney Disease | 88 | 88(0) | 1 |
| Neurodevelopmental Disorders | 52 | 48(4) | 0.923 |
| Hearing Loss | 51 | 50(1) | 0.98 |
| Immunology | 48 | 47(1) | 0.979 |
| Cardiovascular | 40 | 39(1) | 0.975 |
| Inborn Errors of Metabolism | 32 | 32(0) | 1 |
| Neurological Disorders | 17 | 16(1) | 0.941 |
| Ocular | 13 | 13(0) | 1 |
| Skeletal Disorders | 12 | 11(1) | 0.917 |

Table S1: performance of leave-one-phenotype out. Number of variant combinations, True Positive (TP), False Negative (FN) between brackets, and recall for the Random Forest (RF) classifier is reported.

| | Mean | Median | 25th Percentile | 75th Percentile |
|---|---|---|---|---|
| **Specificity** | 0.998 | 0.998 | 0.997 | 0.999 |
| **Recall** | 0.837 | 0.843 | 0.784 | 0.906 |
| **PRC** | 0.915 | 0.925 | 0.891 | 0.950 |
| **Precision** | 0.901 | 0.908 | 0.870 | 0.949 |
| **F1_score** | 0.864 | 0.869 | 0.833 | 0.899 |
| **Balanced Accuracy** | 0.917 | 0.921 | 0.891 | 0.952 |
| **TD recall** | 0.731 | 0.733 | 0.612 | 0.867 |
| **CO recall** | 0.899 | 0.917 | 0.846 | 1.000 |

Table S2. Performance of diVas phenotype-free approach on 500 repeated sampling iterations. Recall is also stratified by True Digenic (TD) and Composite (CO). DMs were not considered for phenotype-free model.

### XAI-based digenic Mechanism prediction

| Attribute | Value | P(DM\|Attribute =value) | P(No-DM\|Attribute=value ) |
|---|---|---|---|
| gene_level_contribution | low | 0.94 | 0.06 |
| | low-median | 0.18 | 0.82 |
| | median-high | 0.03 | 0.97 |
| | high | 0.0 | 1.0 |
| variant_level_contribution | low | 0.09 | 0.91 |
| | low-median | 0.15 | 0.85 |
| | median-high | 0.18 | 0.82 |
| | high | 0.74 | 0.26 |

| | | | |
|---|---|---|---|
| phenotype_level_contribution | low | 0.47 | 0.53 |
| | low-median | 0.26 | 0.74 |
| | median-high | 0.24 | 0.76 |
| | high | 0.18 | 0.82 |

Table S3: Naive Bayes conditional probabilities computed on the training set for each features subgroups contribution to classification.

### Validation on real cases and benchmark analysis

| SampleID | Number of available HPO | Number of analyzed variants | Family availability | Digenic Mechanism |
|---|---|---|---|---|
| Sample1 | 1 | 331 | NO | TD |
| Sample2 | 1 | 144 | NO | TD |
| Sample3 | 3 | 206 | YES | DM |
| Sample4 | 2 | 97 | NO | DM |
| Sample5 | 2 | 121 | NO | NA |
| Sample6 | 1 | 318 | NO | TD |
| Sample7 | 3 | 115 | NO | CO |
| Sample8 | 2 | 349 | YES | DM |
| Sample9 | 1 | 363 | YES | DM |
| Sample10 | 4 | 330 | YES | TD |
| Sample11 | 2 | 296 | YES | TD |

Table S4: General information of the 11 real samples included in the validation.

| Sample ID | Class | | | | | Ranking | | | | | # Patho | | | | |
|---|---|---|---|---|---|---|---|---|---|---|---|---|---|---|---|
| | diVas | ORVAL | DiGePred | diVas Pheno Free | DIEP | diVas | ORVAL | DiGePred | diVas Pheno Free | DIEP | diVas | ORVAL | DiGePred | diVas Pheno Free | DIEP |
| Sample1 | 1 | 1 | 1 | 1 | 0 | 1 | 88(88) | 16 | 3 | NA | 14 | 3454(3235) | 32 | 191 | 463 |
| Sample2 | 1 | 1 | 0 | 1 | 1 | 12 | 159(159) | 3252 | 5 | 346 | 38 | 3364(3364) | 30 | 261 | 551 |
| Sample3 | 1 | 0 | 0 | 0** | 0 | 2 | 4565(4549) | 760 | 31 | 1313 | 8 | 3567(3556) | 8 | 19 | 76 |
| Sample4 | 0 | 1 | 0 | 0** | 0 | 77 | 82(82) | 516 | 189 | NA | 16 | 1037(1037) | 29 | 158 | 219 |
| Sample5 | 0 | 1 | 0 | 1 | 0 | 49 | 182(182) | 467 | 41 | 1558 | 0 | 1774(1774) | 45 | 72 | 453 |
| Sample6 | 1 | NA* | 0 | 1 | 1 | 1 | NA* | 61 | 2 | 133 | 28 | NA* | 45 | 202 | 306 |
| Sample7 | 1 | 1 | 1 | 1 | 1 | 1 | 5(5) | 3 | 1 | 4 | 39 | 807(791) | 25 | 55 | 266 |
| Sample8 | 1 | 1 | 0 | 1** | 0 | 30 | 3758(3674) | 6713 | 86 | 11401 | 36 | 3883(3796) | 62 | 200 | 948 |
| Sample9 | 1 | 1 | 0 | 0** | 1 | 25 | 150(150) | 6016 | 1153 | 385 | 60 | 3771(3624) | 52 | 170 | 662 |
| Sample10 | 0 | 0 | 0 | 1 | 1 | 3 | NA(NA) | 4477 | 1 | 8 | 2 | 4159(3569) | 17 | 32 | 284 |
| Sample11 | 1 | 0 | 0 | 1 | 1 | 1 | NA(NA) | 4669 | 1 | 8 | 2 | 4154(3577) | 17 | 32 | 294 |

Table S5: Benchmark results on 11 real samples analyzed with diVas, ORVAL, DiGePred, the phenotype-free model of diVas and DIEP. For each tool and for each sample, predicted class, ranking and predicted probability of the causative digenic combination is reported. Number of predicted pathogenic combinations is also reported. For ORVAL, we report results calculated both on ORVAL output (with more variant pairs for the same gene pair) and on ORVAL output deduplicated by gene pairs, by selecting for each gene pair the variant combination with the highest predicted probability (results shown in parenthesis).

* ORVAL was not able to show Sample6 results through the web interface, probably due to a high number of analyzed digenic combinations.

** These predictions refer to Dual Molecular Diagnosis. Since diVas phenoFree model was not trained on DMs, we do not expect high performance on this subset.

| SampleID | Digenic Mechanism | diVas predicted digenic mechanism | ORVAL predicted digenic effect |
|---|---|---|---|
| Sample1 | TD | TD/CO | CO |
| Sample2 | TD | DM | TD |
| Sample3 | DM | DM | Neutral |
| Sample4 | DM | Neutral | TD |
| Sample6 | TD | DM | NA |
| Sample7 | CO | TD/CO | CO |
| Sample8 | DM | DM | CO |
| Sample9 | DM | DM | CO |
| Sample10 | TD | Neutral | NA |
| Sample11 | TD | TD/CO | NA |

Table S6: Digenic mechanism predictions obtained with diVas and ORVAL on 10 real samples. TD=True Digenic, CO=Composite, DM=Dual Molecular Diagnosis, NA=Not Available. Neutral is assigned when the causative digenic combination has been classified as benign by the pathogenicity prediction algorithm; NA is assigned when the digenic mechanism prediction was not computed due to missing annotation.

### Figures

#### Monogenic Variant Interpretation

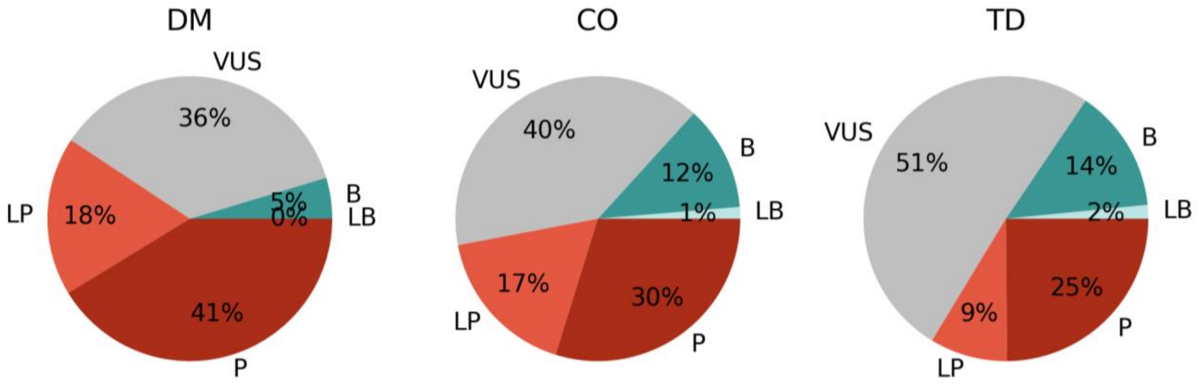


Figure S1: Percentage of variants in DM, CO and TD pairs interpreted as Pathogenic (P), Likely pathogenic (LP), Uncertain (VUS), Likely benign (LB) and Benign (B) according to the ACMG/AMP guidelines.

#### diVas performance on 500 repeated hold-out validation

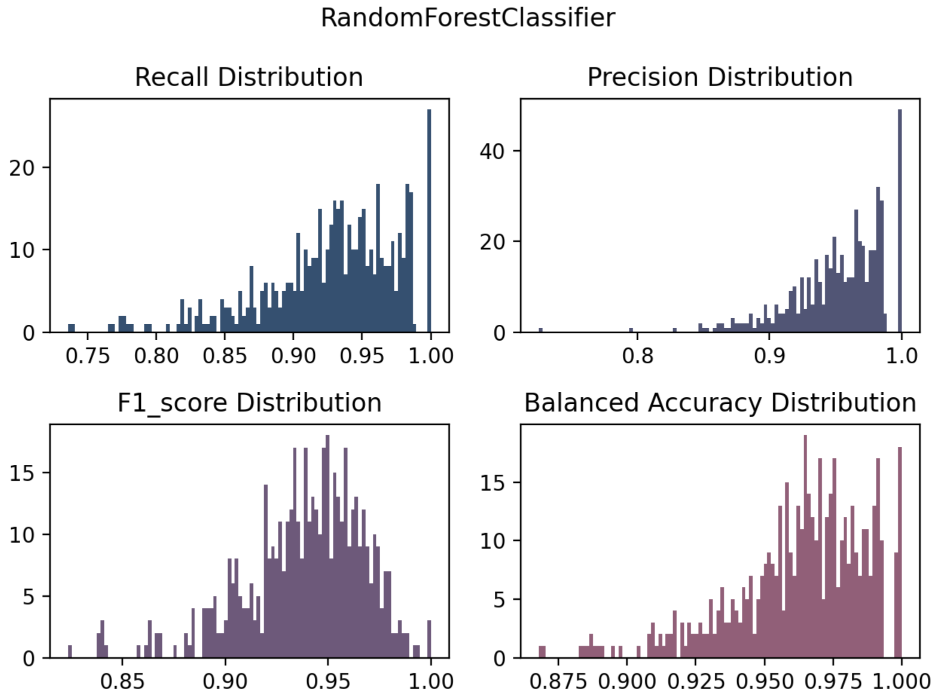


Figure S2: Distribution of Recall, Precision, F1 and Balanced Accuracy computed with 500 repeated sampling for the phenotype-driven RF model.

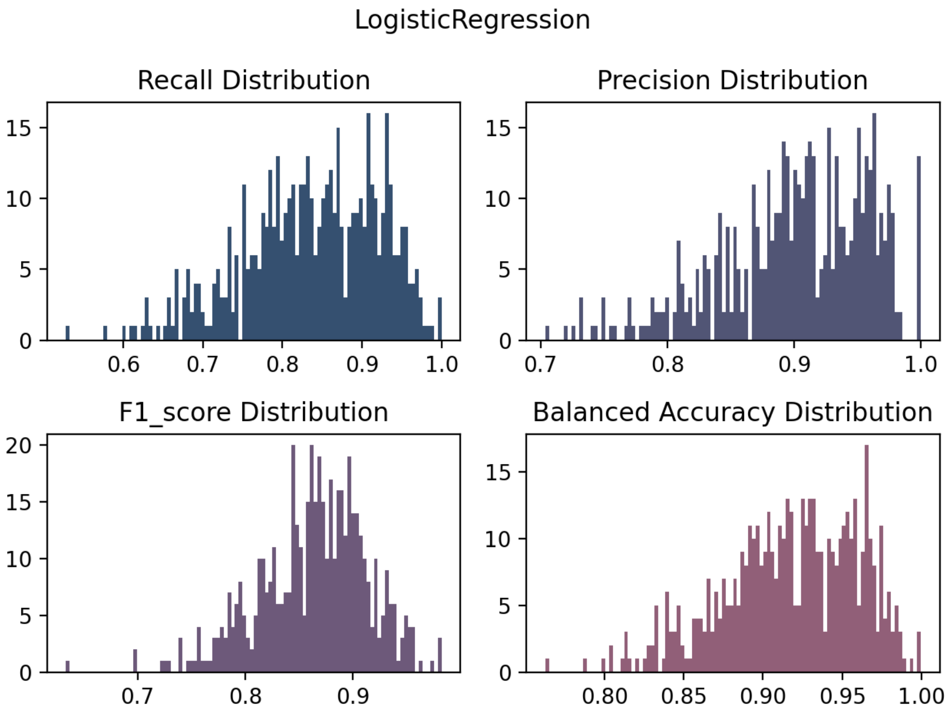


Figure S3: Distribution of Recall, Precision, F1 and Balanced Accuracy computed with 500 repeated sampling for the phenotype-free LR model.
